# Prophages and satellite prophages are widespread among *Streptococcus* species and may play a role in pneumococcal pathogenesis

**DOI:** 10.1101/502740

**Authors:** Reza Rezaei Javan, Elisa Ramos-Sevillano, Asma Akter, Jeremy Brown, Angela B Brueggemann

## Abstract

Prophages (viral genomes integrated within a host bacterial genome) are abundant within the bacterial world and are of interest because they often confer various phenotypic traits to their hosts, such as by encoding genes that increase pathogenicity. Satellite prophages are ‘parasites of parasites’ that rely on the bacterial host and another helper prophage for survival. We analysed >1,300 genomes of 70 different *Streptococcus* species for evidence of prophages and identified nearly 800 prophages and satellite prophages, the majority of which are reported here for the first time. We show that prophages and satellite prophages were widely distributed among streptococci, were two clearly different entities and each possessed a structured population. There was convincing evidence that cross-species transmission of prophages is not uncommon. Furthermore, *Streptococcus pneumoniae* (pneumococcus) is a leading human pathogen worldwide, but the genetic basis for its pathogenicity and virulence is not yet fully understood. Here we report that over one-third of pneumococcal genomes possessed satellite prophages and demonstrate for the first time that a satellite prophage was associated with virulence in a murine model of infection. Overall, our findings demonstrate that prophages are widespread components of *Streptococcus* species and suggest that they play a role in pneumococcal pathogenesis.

## Main

The genus *Streptococcus* comprises a wide variety of pathogens responsible for causing significant morbidity and mortality worldwide^1^. *Streptococcus pneumoniae* (pneumococcus) is a leading cause of pneumonia, bacteraemia, and meningitis^2^. *Streptococcus pyogenes* (group A streptococci) is a leading cause of pharyngitis, scarlet fever and necrotising fasciitis^3^. *Streptococcus agalactiae* (group B streptococci) is the most common cause of neonatal sepsis^4^. *Streptococcus suis* and *Streptococcus equi* rarely cause disease in humans but are important animal pathogens^1^.

Bacteriophages (phages) are intracellular parasites of bacteria. Lytic phages hijack the host bacterial machinery, produce new phages and destroy the infected bacterial cell. Lysogenic phages do not necessarily initiate replication immediately upon host entry and may integrate their genome within the bacterial genome to be activated at a later stage. An integrated phage is termed a prophage and those genes can be passed down to the bacterial daughter cells.

Since survival depends on their bacterial hosts, prophages often express genes that increase host cell fitness^5,6^. Prophages can exert a range of phenotypic effects on the host bacteria: encode toxins that increase virulence^5^, promote binding to human platelets^7^ or cells^8^, evade immune defences^9,10^, or protect from oxidative stress^11^. Prophage integration can also regulate bacterial populations by altering bacterial gene expression^12,13^.

Prophages and their hosts, like other predator and prey relationships, are embroiled in a complex evolutionary arms race whereby bacteria evolve various strategies to defend themselves and prophages co-evolve to overcome these barriers^14^. These coevolutionary dynamics are complicated by satellite prophages, which lack all the necessary genetic information to replicate on their own and are reliant on hijacking the machinery of another inducing ‘helper’ prophage to replicate. Satellite prophages might be thought of as ‘parasites of parasites’^15,16^.

Satellite prophages adversely interfere with helper prophage replication and thus promote bacterial survival^17-19^. Satellite prophages have been discovered through different circumstances and thus there are different terms used to describe this particular type of mobile genetic element (MGE) in the literature, including *Staphylococcus aureus* pathogenicity islands (SaPIs), phage-related chromosomal islands (PRCIs) and phage-inducible chromosomal islands (PICIs), among others^17,23^.

Satellite prophages have been shown to be vectors for the spreading of toxin genes and other virulence factors, e.g. SaPI1, which possesses the gene responsible for causing toxic shock syndrome^24^. The prevalence, diversity, genetic stability and molecular epidemiology of satellite prophages in streptococcal species are largely unknown. A small number of satellite prophages have been identified in streptococcal species, although whether they are associated with virulence remains to be investigated^25^. Previous work has shown that prophage-related sequences are highly prevalent within *S*. *pneumoniae*^26-28^, *S*. *pyogenes^25,30^* and *S*. *agalactiae* genomes^31^; however, genus-wide analyses of the genomic diversity and population structure of streptococcal prophages have not yet been reported.

Here we report the discovery of nearly 800 prophages among >1,300 streptococcal genomes and provide detailed insights into prophage genomics and population structure. Using *S*. *pneumoniae* as the model organism, the molecular epidemiology of satellite prophages was investigated within a large globally-distributed collection of pneumococci isolated over a 90-year period. Finally, we demonstrated that a satellite prophage was associated with virulence in a murine infection model.

## Results

### Prophage sequences are a significant component of the genomes of clinically-relevant *Streptococcus* species

We analysed 1,306 genomes from 70 different streptococcal species and identified 419 full-length prophages and 354 satellite prophage genomes (Supplementary Table 1). We estimated the prophage gene content within each streptococcal genome and this revealed a substantial difference in the average prophage content among various streptococcal species, ranging from 0.4% of the *Streptococcus thermophilus* genome to 9.5% of the *S*. *pyogenes* genome (Figure 1a; Supplementary Table 2). Furthermore, we observed significant variability in prophage content among different genomes of the same bacterial species, e.g. full-length prophages comprised up to 19% of the genes in some *S*. *pyogenes* genomes, while in others they made up <1% of the genome (Figure 1a). The prevalence of satellite prophages ranged from 0.1% among *Streptococcus mutans* and *Streptococcus sanguinis* genomes to 4.5% of the *Streptococcus dysgalactiae* genomes (Figure 1a).

**Fig. 1.**
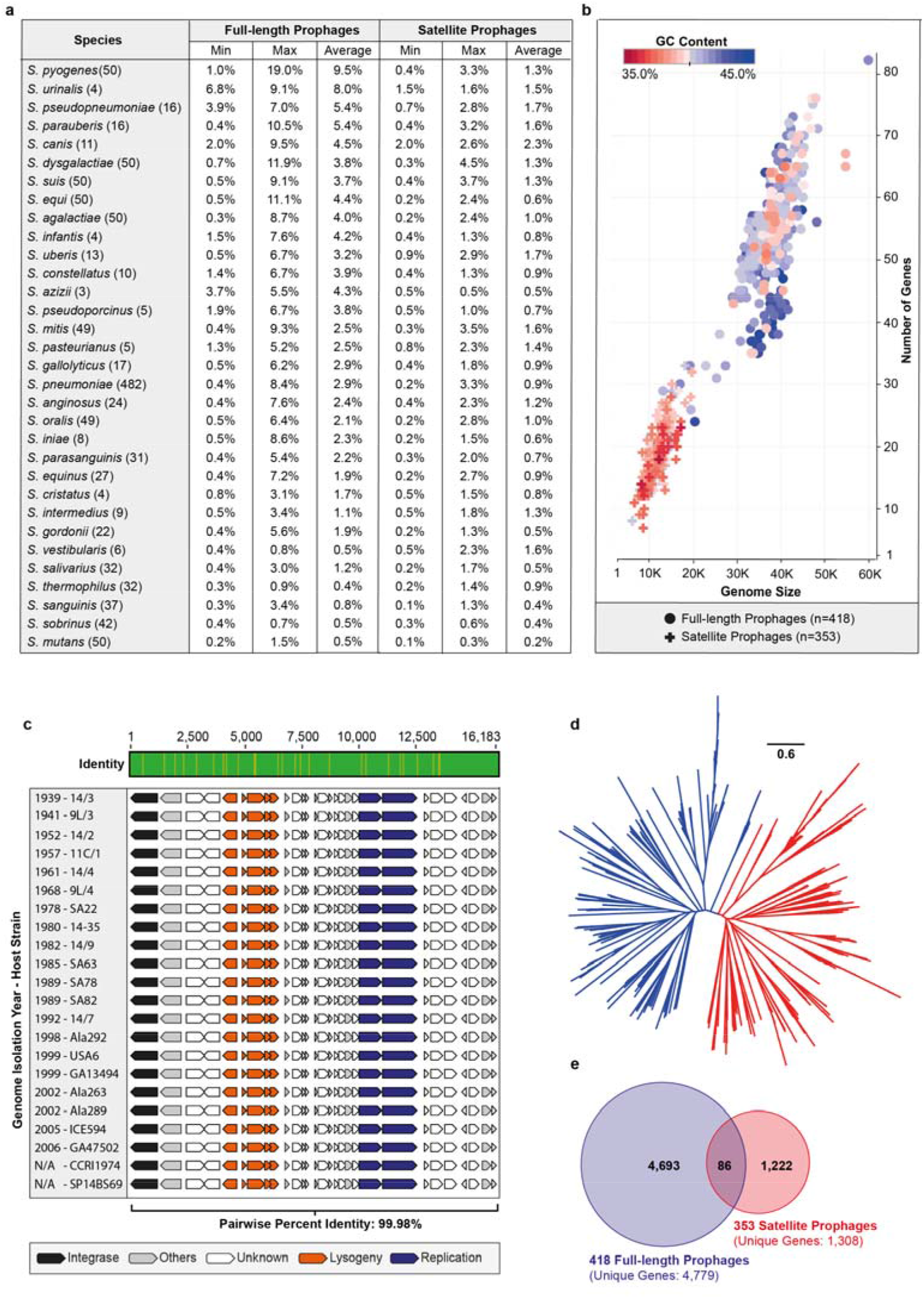
Full-length and satellite prophages identified among streptococcal genomes. **a**, Average prophage content within each streptococcal species. **b**, Graphical representation of all prophages by average genome size and number of genes. Each prophage is coloured to represent its average guanine (G) and cytosine (C) content. **c**, Satellite prophage SpnSP24 was represented among pneumococci isolated between 1939 and 2006 and all of these satellite prophages were nearly identical at the nucleotide level one to another. **d**, An unrooted phylogenetic tree of all streptococcal prophage genomes identified in the dataset. Blue branches mark full-length prophages and red branches mark satellite prophages. **e**, Genes found in full-length prophages and in satellite prophages (at a threshold of >70% amino acid sequence similarity) are rarely shared.

### Full-length and satellite prophages are separate entities with little effective genetic exchange between them

Satellite prophages had a lower guanine (G) and cytosine (C) content than full-length prophages and were about a third of the size in terms of both length of sequence and the number of genes they harboured (Figure 1b). Due to their relatively small genome and apparent lack of essential genes, streptococcal satellite prophage sequences have historically often been regarded as “remnant” or “defective” prophages in a state of mutational decay^13,22,32-34^. Our data reveal that satellite prophage sequences can be highly conserved over many decades, e.g. one satellite prophage was present among pneumococcal genomes with isolation dates ranging from 1939 to 2006 and had maintained >99.98% nucleotide similarity across its entire genome (Figure 1c), suggesting that it is under very strong evolutionary pressure and likely provides an important biological function.

An unrooted phylogenetic tree of all streptococcal prophage genomes in our dataset depicted full-length and satellite prophages as two clearly distinct groups (Figure 1d). We observed that the genes of satellite prophages are unique and differ to those of full-length prophages, as nearly 99% of all satellite prophage genes (>70% amino acid sequence similarity) are not found in any full-length prophages (Figure 1e). Taken together, these findings confirm that satellite prophage sequences are not recent remnants of previous lysogenisation by full-length prophages, but rather that they belong to a unique family of mobile genetic elements.

### Streptococcal prophages have a structured population

We found that both full-length and satellite streptococcal prophages demonstrated well-conserved patterns in genome organisation and synteny, regardless of the species that they were isolated from (Figure 2a). Similar to other non-streptococcal prophages (Supplementary Figure 1), genes encoding specific functions were often found clustered together in the prophage genome, although note that the function of many genes is still unknown and therefore the delineation of discrete gene clusters remains problematic (Figure 2a). Whole genome comparisons of all prophage sequences in our dataset depicted several major and minor clusters for both full-length and satellite prophages (Supplementary Figure 2).

**Fig. 2.**
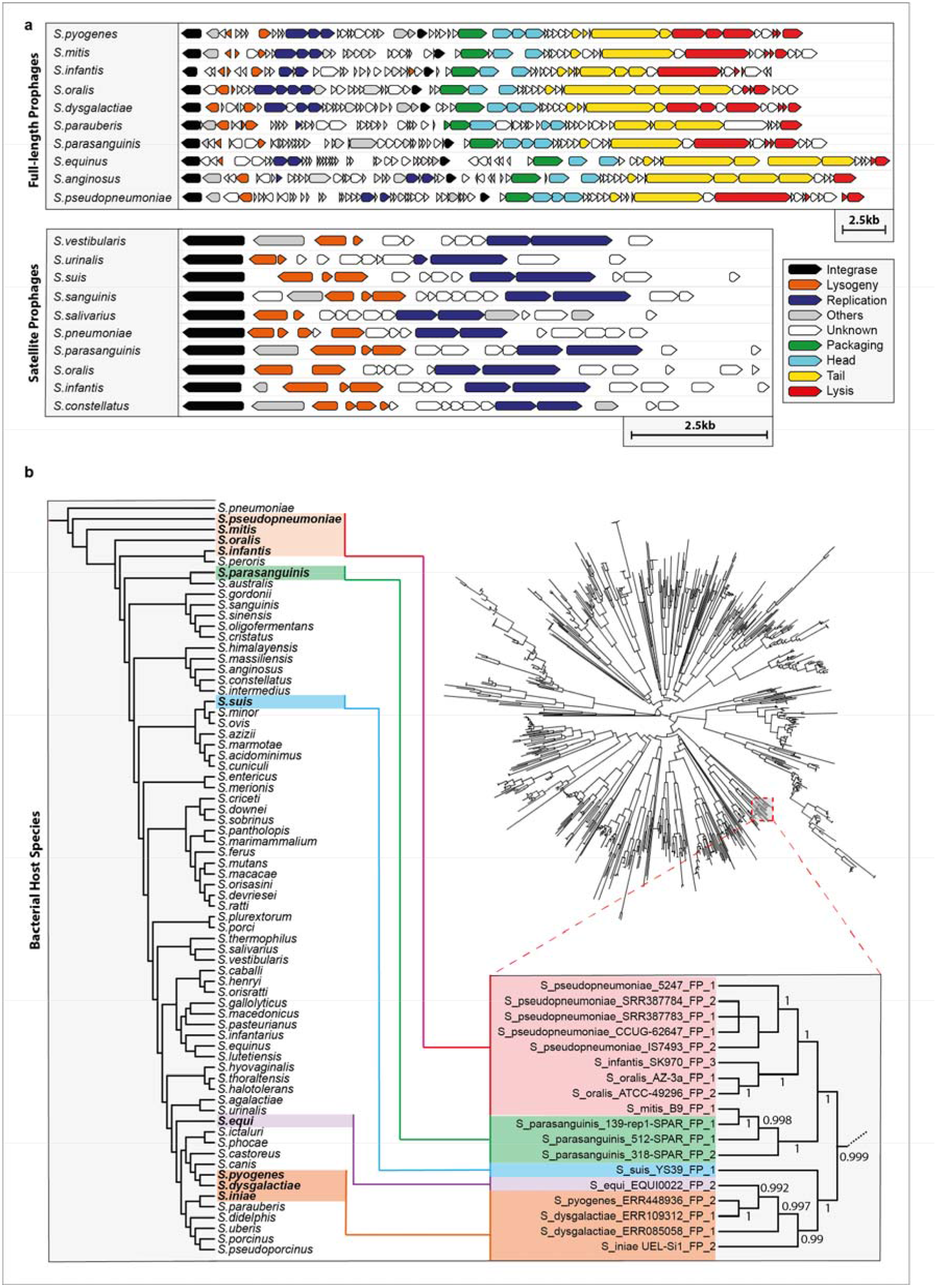
Similarities among streptococcal prophages and evidence for cross-species transmission of prophages. **a**, Full-length and satellite prophages identified among different streptococcal species shared an identical pattern in gene orientation and synteny. **b**, Phylogenetic tree depicting all prophages detected in this study (see Supplementary Fig. 2 for a larger version of the tree) and a zoomed-in branch depicting one example of a cluster of full-length prophages that were found among multiple streptococcal species.

Phages are generally believed to be bacterial species-specific and even specific to genetic lineages within a single bacterial species^35^. Surprisingly, we often found prophages from different bacterial species within the same phylogenetic cluster, suggesting that cross-species transmissions are more common among streptococcal prophages than previously realised. Remarkably, despite the relatedness of their prophages, the bacterial hosts were not necessarily the closest phylogenetically-related species (Figure 2b). One possible explanation could be that streptococcal prophages are evolving separately from their microbial hosts, and therefore, other factors such as ecological relatedness may dominate over evolutionary relatedness of the host bacteria.

### Molecular epidemiology of satellite prophages within a global pneumococcal dataset dating from 1916

We had previously determined the prevalence, diversity and molecular epidemiology of full-length prophages in a global and historical pneumococcal genome dataset^26^. Many shorter prophage sequences were also identified in that study, which were simply classified as partial prophage sequences and not characterised further at the time. Here, we used this genome dataset to further investigate satellite prophages in the context of the pneumococcal population structure. The genome collection was comprised of 482 pneumococci recovered from both healthy and diseased individuals between 1916 and 2009. Pneumococci were isolated from people of all ages residing in 36 different countries. Ninety-one serotypes and 94 different clonal complexes (genetic lineages) were represented in the dataset.

A reinvestigation of the ‘partial prophage’ sequences resulted in the identification of 44 representative pneumococcal satellite prophages, which clustered into five major groups (Figure 3a). The average GC content of the satellite prophages was lower than their pneumococcal host but varied among each group (Figure 3b). We found that 35% of the pneumococci in our dataset contained at least one satellite prophage and 5% of the genomes contained two. Some satellite prophages were present in up to six different clonal complexes, whereas others were only found in Singletons (genotypes with no closely related variants; Table 1 and Supplementary Figure 3). Those satellite prophages identified in more than one genome were often found among pneumococci recovered over a decade or more (Table 1). The average prophage content for each of the major clonal complexes ranged from 2.2–6.5%, and with only one exception (CC7232), all of these are widely-circulating pneumococcal genetic lineages (Figure 1c; https://pubmlst.org/spneumoniae).

**Fig. 3.**
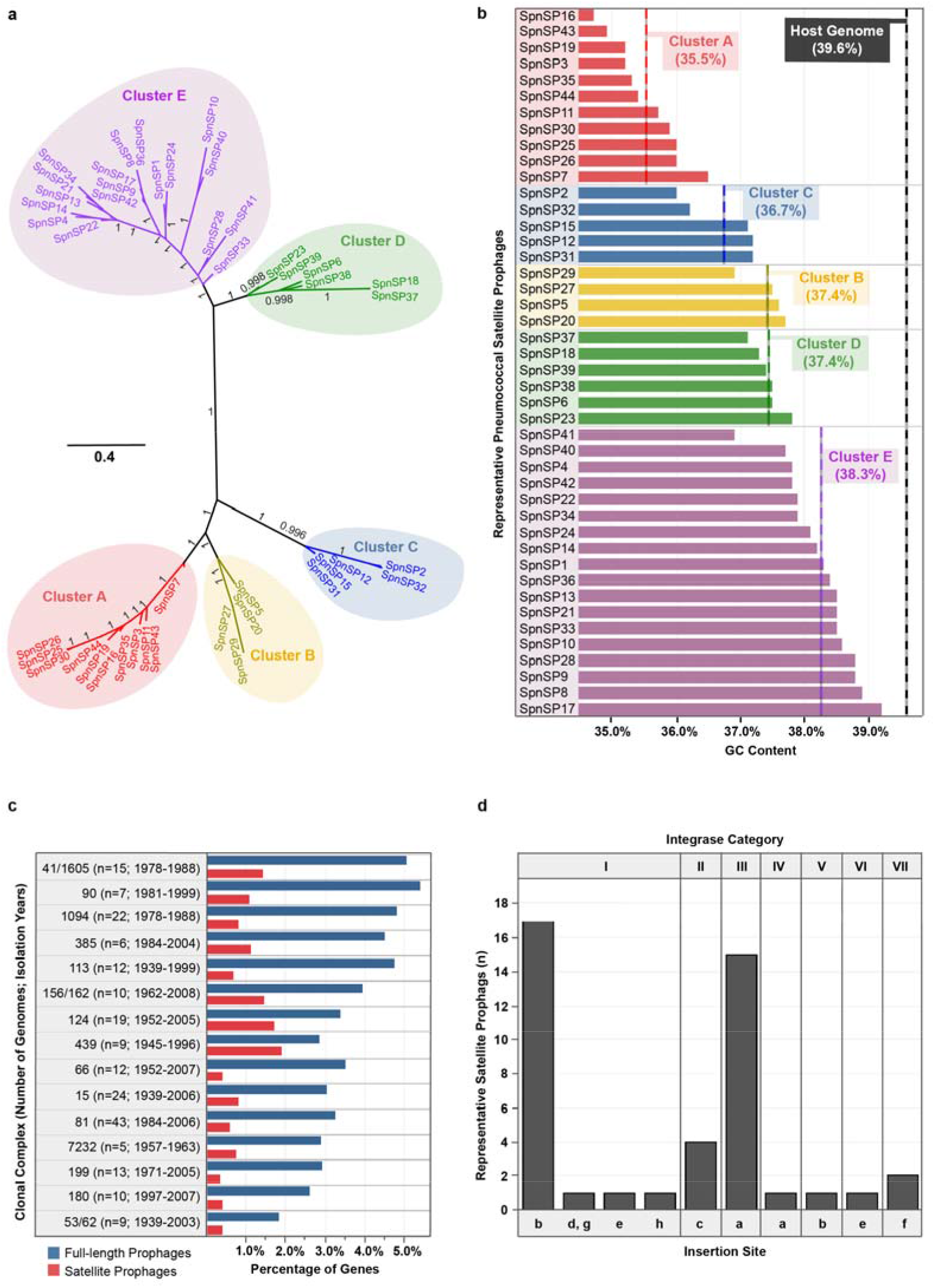
Satellite prophages found among a large collection of nearly 500 diverse pneumococcal genomes. **a**, An unrooted phylogenetic tree demonstrated that the 44 representative satellite prophages could be clustered into five major groups based upon nucleotide similarity. b, The average guanine/cytosine (GC) content (stated in brackets) of the satellite prophages varied by genetic cluster and was lower than the GC content of the pneumococcal host. **c**, The average prophage content for each of the major clonal complexes (genetic lineages) is depicted as a percentage of the total number of genes in the host pneumococcal genome (~2 Mb). **d**, The integrase sequences of the 44 representative satellite prophages were divided into seven different categories based upon ≥95% nucleotide similarity.

**Table 1.**
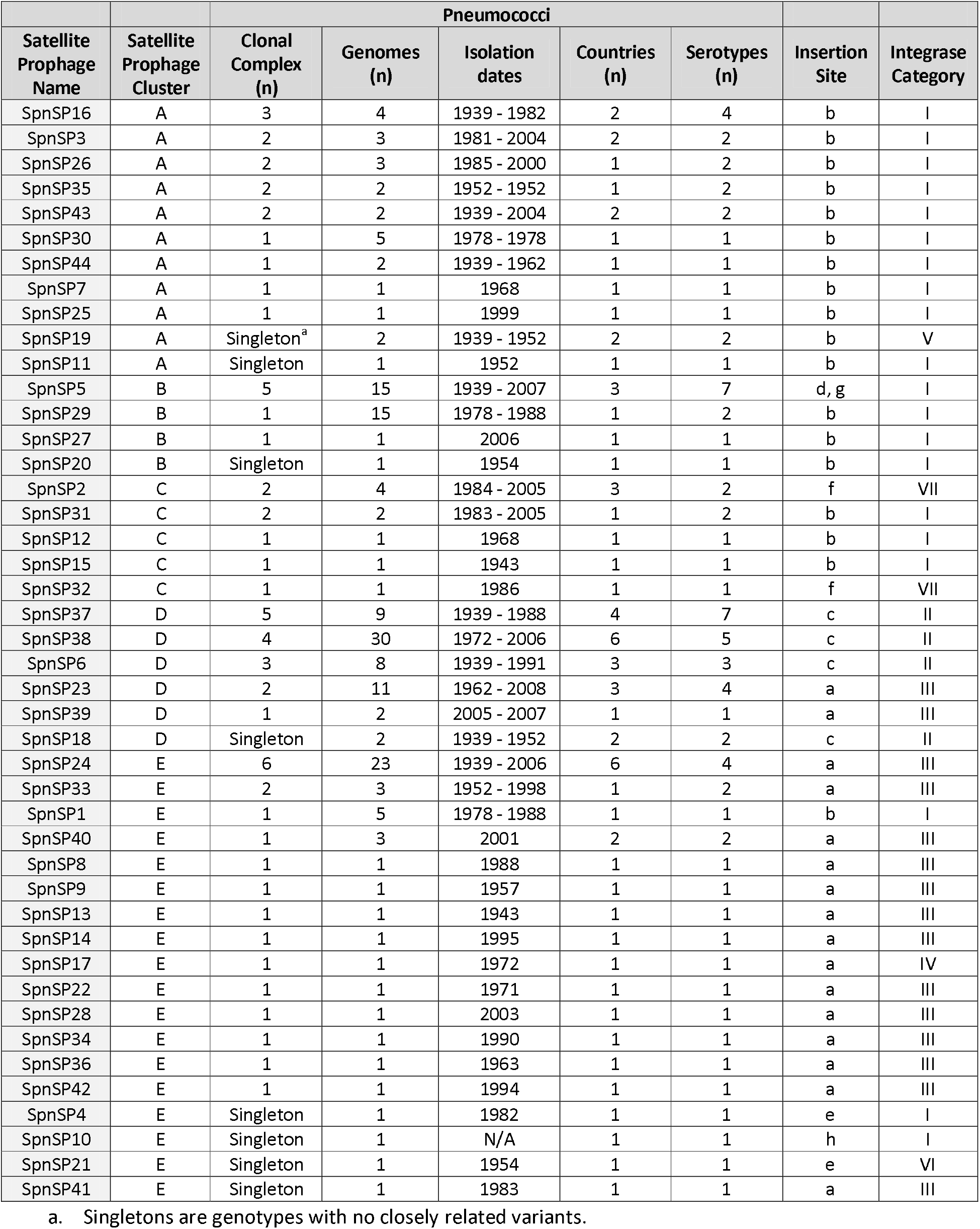
Epidemiological characteristics of 44 representative satellite prophages identified among a collection of pneumococcal isolates dating from 1939 onwards.

### Prophages are more frequently inserted adjacent to genes involved in information storage and processing

We previously reported that pneumococcal full-length prophages were consistently integrated in specific locations within the genome^26^. Likewise, pneumococcal satellite prophages were consistently integrated in seven precise locations (a-f) within the host genome, each of which was directly associated with the integrase gene they harboured (Figure 3d; Figure 4a). The 44 representative satellite prophage integrases were divided into seven different categories with ≥95% nucleotide sequence similarity within each category. Each integrase category was associated with insertion at a single location on the pneumococcal genome, apart from integrase category I, which was associated with five different locations (Figure 3d). 28.3% of pneumococcal satellite prophages were inserted at site a, which was very close to the origin of replication (oriC) (Figure 4a) and prompted us to investigate whether factors other than the integrase sequence determined the prophage insertion site.

**Fig. 4.**
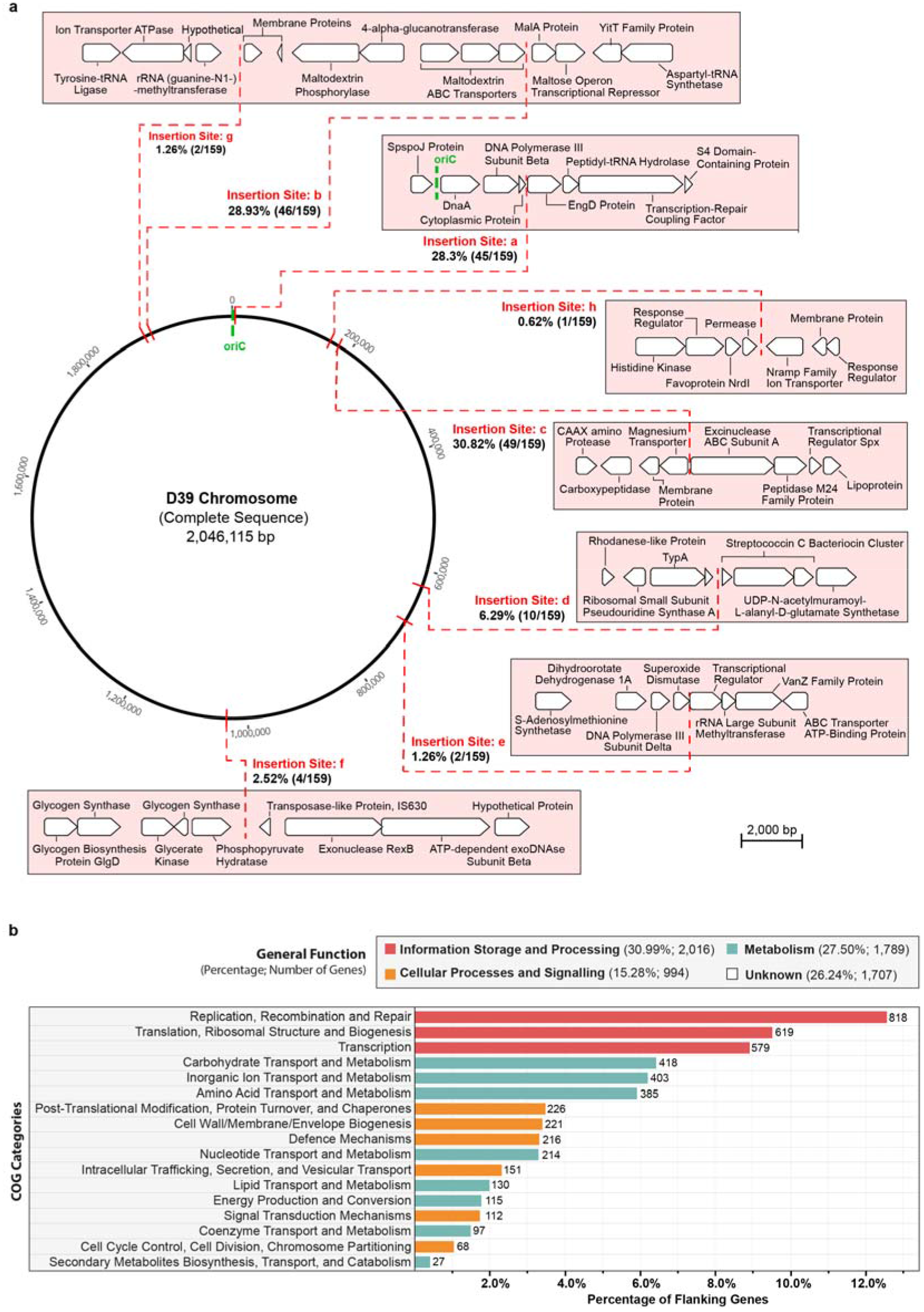
Insertion sites of prophages within the pneumococcal genome. **a**, Pneumococcal satellite prophages were integrated in seven locations (a-f) within the host genome. Percentages and numbers in brackets refer to the proportion and number out of all 159 satellite prophages that were inserted in that particular location. **b**, The pneumococcal flanking genes upstream and downstream of all integrated full-length and satellite prophages were retrieved for functional classification and are depicted here based upon their COG (clusters of orthologous groups) classifications.

We investigated the location of prophage insertion sites within the genome sequences of non-pneumococcal streptococci for which at least one complete genome was available (n=29). We divided the genome of each species into 8 non-overlapping segments of equal length according to the number of base pairs, and the percentages of prophages situated in each segment were quantified. Overall, we observed no strong preference for prophage insertion in any of the 8 segments and the location of prophages residing within the genome varied greatly between different species (Supplementary Figure 4).

Among pneumococcal and non-pneumococcal streptococcal genomes, five flanking genes upstream and downstream of each prophage were retrieved for functional classification using gene ontology analyses. This revealed that nearly one-third of all the bacterial flanking genes were involved in replication, recombination, DNA repair, transcription, translation and ribosomal structure and biogenesis (Figure 4b). One-quarter of flanking genes were involved in metabolic processes, but equally, one-quarter of all flanking genes did not have a defined functional classification. The remaining flanking genes were involved in other cellular processes and signalling. A list of all prophage insertion sites and their flanking genes is available in Supplementary Table 3.

### Satellite prophages and *vapE* are involved in pneumococcal pneumonia and sepsis in a murine infection model

Our investigation of pneumococcal satellite prophage genes led to the identification of a gene that is a homologue of the ‘virulence-associated gene E’ *(vapE)* in *S*. *suis^36^*. We investigated *vapE* in *S*. *suis* genomes and confirmed that it is carried by a satellite prophage. We searched for *vapE* in the representative pneumococcal satellite prophages and found that 30/44 (68.18%) contained *vapE*. To investigate whether the *vapE* homologue in the pneumococcal satellite prophage is also associated with virulence, we performed *in vivo* studies using a murine pneumococcal infection model.

Deletion mutant stains were constructed in a serotype 6B pneumococcal strain (BHN418) in which either *vapE* only *(ΔvapE)* or the entire satellite prophage sequence *(ΔSpnSP38)* were replaced by a spectinomycin resistance cassette *(aadA9)*. Both mutant pneumococcal strains grew as well as the parental wild-type strain in Todd-Hewitt plus yeast extract broth media (data not shown). For each of the mutant strains a competitive index (CI) was determined using a highly sensitive competitive infection experiment in a mouse model of pneumonia.

The CI was significantly lower in the lungs after mixed infection with *ΔSpnSP38* vs serotype 6B or *ΔvapE* vs serotype 6B, indicating a role for the satellite prophage and *vapE* in the establishment of pneumococcal pneumonia (Figure 5a). To further assess the degree of attenuation in virulence of the *ΔSpnSP38* and *ΔvapE* strains, infection experiments were repeated with pure inocula of each strain in both the pneumonia and sepsis models. There were no significant differences in bacterial CFU recovered from the lungs of infected mice at 24 h between either mutant and the parental wild-type strain (Figure 5b) and the majority of the mice developed fatal infection by this point. However, in the sepsis model the mice infected with the wild-type serotype 6B strain had significantly greater blood and spleen CFU than the *ΔSpnSP38* mutant (Figure 5c and 5d), indicating that the satellite prophage is directly involved in pneumococcal virulence during bacterial dissemination in the systemic circulation. Although the *ΔvapE* strain had lower spleen CFU compared to the wild-type, this difference was not statistically significant, suggesting that loss of the whole satellite prophage has a more marked effect on the attenuation of virulence during sepsis than loss of VapE alone.

**Fig. 5.**
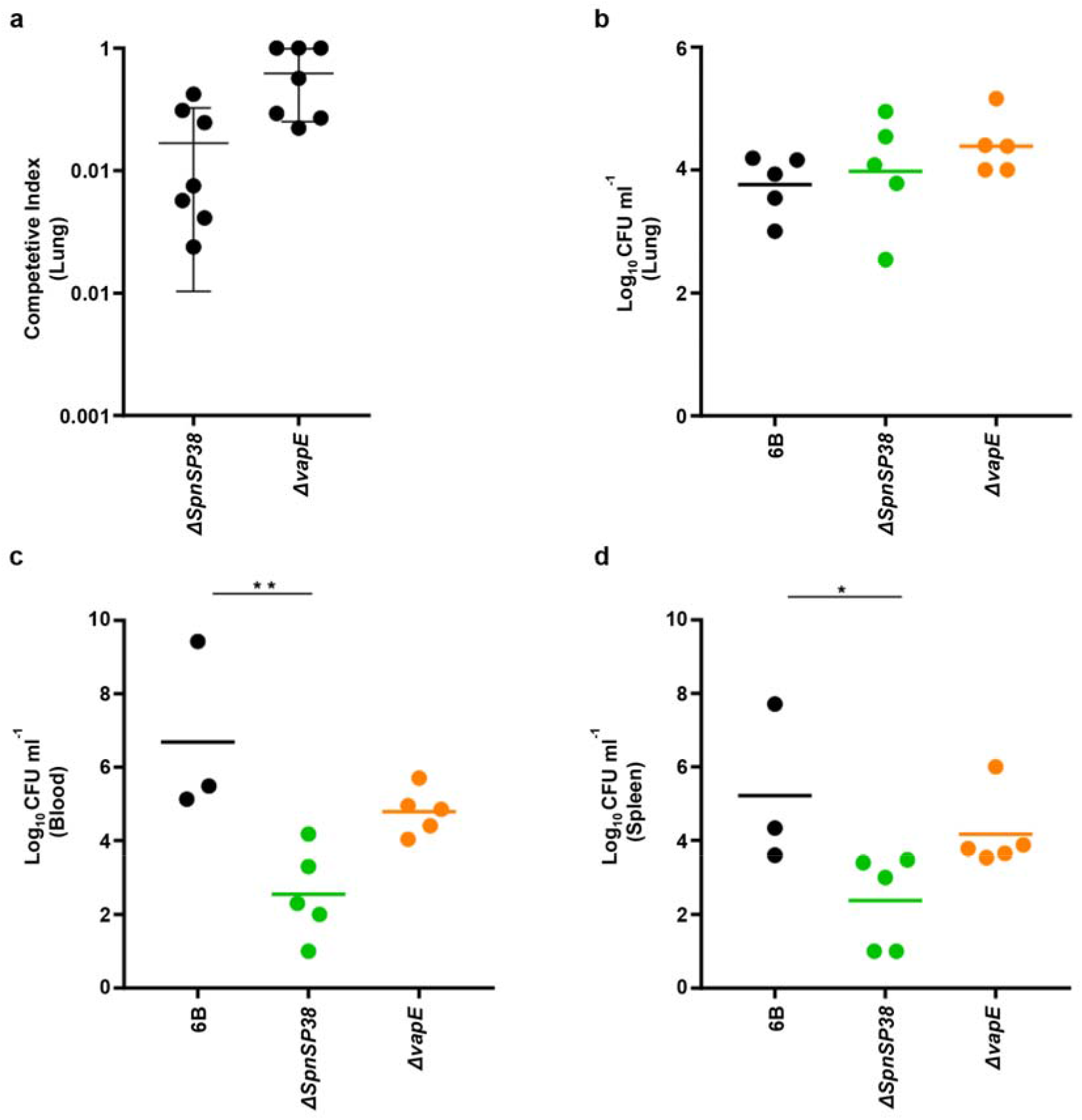
Assessment of the virulence of *ΔSpnSP38* (deletion of entire satellite prophage) and *ΔvapE* (deletion of *vapE* only) mutant pneumococcal strains in murine infection. **a**, A comparison of the competitive index (CI) in the lungs in a model of pneumonia of mixed infection with the parental wild-type strain versus the mutant. Each point represents the CI for a single animal. **b**, Bacterial levels recovered from lung homogenates at 24h (expressed as colony-forming units (CFU) per ml) after pneumococcal pneumonia produced with the wild-type and mutant strains. **c, d**, Bacterial levels recovered at 28h from blood or spleen homogenate. Error bars represent standard error of the mean (Kruskall-Wallis with Dunn’s post hoc test)<ENT FONT=(normal text) VALUE=46>.</ENT>

## Discussion

In this study we revealed an extraordinarily diverse collection of full-length prophages and satellite prophages among streptococcal species. What was striking about these findings was that prophages and satellite prophages were two clearly different entities and both had a structured population. Specifically, among pneumococci there were prophages with persistent associations to genetic lineages of pneumococci over long periods of time. This is crucial, since these data allow for the exploration of *why* certain combinations of prophages and bacteria exist and whether the prophages might be contributing to the epidemiological success of bacterial genetic lineages.

Our findings suggest that prophages are likely to be influencing bacterial biology and epidemiology to a much greater extent than previously appreciated, given the high proportion of prophage DNA present in many streptococcal species. Prophages are mobile genetic elements and genetically similar prophages were frequently detected between different streptococcal species. This implies that prophage transmission across bacterial species is more common than previously recognised, which should be taken into account when trying to understand the precise role of prophages in streptococcal biology.

Many of the streptococci we investigated are important human and animal pathogens and we demonstrated that a previously unrecognised pneumococcal satellite prophage was significantly associated with virulence in a murine model of infection. The mechanism driving virulence is not yet clear, but this work is proof-of-principle that experimental investigations of pneumococcal prophages should be pursued, and these may reveal central aspects of the bacteria/prophage relationship among pneumococci and other streptococci.

The increasingly large volume of genome sequence data in the public domain presents many new opportunities for understanding bacterial infection and pathogenesis at a depth and breadth never before experienced. Large population-level analyses such as this alter our perspective on how bacterial and prophage populations interact and drive evolution of both parasite and host. As demonstrated here, population genomics studies can and should be used to generate hypotheses, design experiments and select the most appropriate strains for testing. The findings of this study reveal numerous areas for further investigation, the results of which will increase our knowledge of prophage and bacterial biology, epidemiology and evolution.

## Methods

### Development of PhageMiner, a bioinformatics tool for prophage identification in bacterial genomes

Some *in silico* prophage detection tools are available that identify prophages using a reference database of known prophage genomes, thus their performance is strongly influenced by the size and composition of the reference dataset^37,38^. In order to ensure a thorough discovery of previously unidentified prophages, manual curation of annotated genomes is required, however, this is not feasible for large genome studies^26,39–40^. To address these issues, we developed a user-supervised semi-automated computational tool called PhageMiner in order to streamline the otherwise tedious manual curation process for prophage sequence discovery. PhageMiner uses a mean shift algorithm combined with annotation-based genome mining in order to rapidly identify prophage sequences within complete or draft bacterial genomes. The source code of PhageMiner is deposited in GitHub (pending).

### Genomes used in this study

In total, 1,316 assembled genomes from 70 different species of the genus *Streptococcus* were selected for this study. 482 genomes belonged to a pneumococcal dataset previously characterised by us^26^. This collection was designed to be highly diverse and consisted of pneumococci recovered from both ill and healthy individuals of all ages residing in 36 different countries between 1916 and 2009. These pneumococci represented 91 serotypes and 94 different clonal complexes (Supplementary Table 4).

The remaining 834 streptococcal genomes were selected from a non-pneumococcal *Streptococcus* species genome dataset previously compiled by us^41^. In brief, 69 different *Streptococcus* species were included in this dataset and up to 50 genomes per species were selected for analyses from the ribosomal MLST database (https://pubmlst.org/rmlst)^42^. When more than 50 genomes were available, the population structure of the species was depicted using PHYLOViZ^43^ and genomes were selected to maximise the population-level diversity of the species from the available genomes. All streptococcal genome sequences were stored in a BIGSdb database^44^ and annotated using the RAST server (http://rast.nmpdr.org).

### Sequence analyses of prophages

All putative prophage sequences were inspected manually using Geneious version 11.1 (Biomatters Ltd.) and those containing ambiguous bases (N’s) and/or assembly gaps were excluded from further analyses. The total number of open reading frames (ORFs), overall sequence length and GC content of each prophage were calculated within the Geneious environment. All multiple sequence alignments were performed using ClustalW^45^ with default parameters (Gap open cost = 15, Gap extend cost = 6.66). Phylogenetic trees were constructed based upon sequence alignments using FastTreeMP^46^. Unique integrase sequences were identified using the CD-HIT program^47^ and a threshold of ≥95% sequence identity. Schematic diagrams of the coding regions of the prophages were produced in Geneious and edited using Adobe Illustrator. The phage content was estimated based on the percentage of prophage genes within a given bacterial genome. To do this, we developed a Python script that first used Prodigal software in the Prokka annotation suite^48^ to predict ORFs in three separate groups of sequences: (i) all identified full-length prophage genomes, (ii) all identified satellite prophage genomes and (iii) a single bacterial genome of interest for which the phage content is to be estimated. Next, the individual ORF nucleotide sequences from all three groups were extracted, combined and clustered using Roary^49^ set at a 70% similarity threshold. Any ORFs in the bacterial genome that were also present in at least one prophage genome were deemed to be phage-related, and this information was used to output the total percentage of phage-related ORFs in the given bacterial genome. The PhageContentCalculator script is available in GitHub (pending).

### Investigation of prophage insertion sites

The prophage insertion sites within the bacterial genome sequences were investigated among the representative pneumococcal prophages and any streptococcal species for which at least one complete bacterial genome was available. To investigate the bacterial genes flanking the prophage sequences, five genes both upstream and downstream of each prophage were retrieved and functional annotations were determined using eggNOG-mapper^50^ based on eggNOG 4.5 orthology data^51^. Prophage insertion sites containing ambiguous bases or assembly gaps were excluded from the analyses. Bacterial genomes were divided into 8 equally-sized segments and the prevalence of prophages per segment was calculated using an in-house Python script (available upon request).

### Construction of a pneumococcal core genome phylogenetic tree

The 482 pneumococcal genomes in the study dataset were annotated using Prokka in order to create GFF3 files compatible with downstream analysis scripts. Genes present in all strains were clustered at 90% sequence identity threshold and aligned using Roary. The phylogenetic tree was generated using FastTreeMP^46^ using a generalized time-reversible model and then was reconstructed using ClonalFrameML^52^ to account for recombination. The tree was annotated using iTOL^53^ and Adobe Illustrator (Adobe Inc.).

### Estimate of phylogenetic relationships among *Streptococcus species*

A phylogenetic tree was constructed using concatenated sequence data from 53 ribosomal loci among all streptococcal genomes in the study dataset using the BIGSdb PhyloTree plugin. The tree was graphically simplified to the species level by collapsing clades containing genomes from the same species into a single leaf using iTOL.

### Bacterial strains, media and growth conditions

*S*. *pneumoniae* strains were cultured in the presence of 5% CO_2_ at 37°C on Columbia agar (Oxoid) supplemented with 5% horse blood, or in Todd-Hewitt broth supplemented with 0.5% yeast-extract (THY; Oxoid). Mutant strains were selected by using appropriate antibiotics (150 μg/ml spectinomycin). Growth of *S*. *pneumoniae* strains in broth was monitored by measuring optical density at 580 nm (OD_580_) and stocks of *S*. *pneumoniae* were stored as single use 0.5 ml aliquots of THY broth culture (OD_580_ 0.4–0.5) at −70°C in 10% glycerol.

### Construction of *ΔvapE* and *ΔSpnSP38 S. pneumoniae* mutant strains

Strains, plasmids and primers used for this study are described in Supplementary Table 5. Both mutants, *ΔvapE* and *ΔSpnSP38* were generated by overlap extension PCR^54,55^ in the *S*. *pneumoniae* serotype 6B BHN418 strain using a transformation fragment in which the *Spn_00749* gene *(vapE)* or the entire satellite prophage, *Spn_00738-Spn_00753*, were replaced by the spectinomycin resistance cassette *aadA9*. For the satellite prophage, two products corresponding to 762 bp upstream (primers SpnSP_UpF and SpnSP_UpspecR) and 872 bp downstream (primers SpnSP_Downspec_F and SpnSP_DownR) of the satellite prophage were amplified from *S*. *pneumoniae* genomic DNA by PCR carrying 3’ and 5’ linkers complementary to the 5’ and 3’ portion of the *aacA9* gene respectively. *aadA9* was amplified from pR412 plasmid (a gift from M. Domenech) using PCR and primers SpnSP_Upspec_F and SpnSP_Downspec_R^54^.

Similarly, for the in-frame deletion of *vapE*, a construct was created in which 820 bp of flanking DNA upstream of the *vapE* ATG (primers VapE_UpF and VapE_UpspcR) and 526 bp of flanking DNA downstream from the *vapE* ORF (starting from the ATG of the overlapping Spn_00750 ORF, primers VapE_DownspecF and VapE_DownR) were amplified by PCR and fused with the *aadA9* cassette by overlap extension PCR^56^. The resulting constructs were then transformed into the BHN418 strain by homologous recombination and allelic replacement using a mix of CSP-1 and CSP-2 and standard protocols^57,58^. The mutations were confirmed by PCR analysis and sequencing.

### Experimental models of infection

6-week-old female CD-1 mice were obtained from Charles River Laboratory and bred in a conventional animal facility at University College of London. Animal procedures were performed according to United Kingdom (UK) national guidelines for animal use and care and approved by the UCL Biological Services Ethical Committee and the UK Home Office (Project Licence PPL70/6510). Studies investigating pneumococcal sepsis or pneumonia were performed using 6-week-old mice and infected as previously described^59^.

Briefly, in the sepsis model mice were challenged with 5 × 10^6^ CFU/ml of the serotype 6B strain or the correspondent mutants in a volume of 150 μl by the intraperitoneal route, whereas for pneumonia, mice under anaesthesia with isofluorane were inoculated intranasally with 50 μl containing 10^7^ CFU/mouse of the serotype 6B strain or the mutants. A lethal dose of pentobarbital was administered at 24 or 28 h after challenge and bacterial counts were determined from samples recovered from lung and blood. Lungs and spleens were homogenised through a 0.2 μm filter. Results were expressed as log_10_ CFU/ml of bacteria recovered from the different sites.

For mixed infection experiments, mice were inoculated with a 50/50 mixture of wild-type and mutant *S*. *pneumoniae*. The competitive index (CI) was defined as the ratio of the test strain (mutant strain) compared to the control strain (wild-type strain) recovered from mice divided by the ratio of the test strain to the control strain in the inoculum^60,61^. A CI of <1 indicates that the test strain is attenuated in virulence compared to the control strain and the lower the CI the more attenuated the strain. Statistical analyses were performed using analysis of variance (ANOVA) for multiple comparisons. GraphPad Prism 7.0 (GraphPad Software, San Diego, CA) was used for statistical analysis.

## Data availability

The newly discovered full-length and satellite prophage genomes have been deposited at GenBank (accession numbers pending). The nucleotide sequence of the *vapE* gene is available via GenBank accession number (pending). Accession numbers for all genomes used in this study are listed in the Supplementary Table 3.

## Supporting information

Supplemental_FigTab_6of9

## Author contributions

R.R.J. and A.B.B. conceived and designed the overall study. R.R.J. wrote the computer code. E.R.S. and J.B. designed the pneumococcal mutants and animal experiments. E.R.S. and A.A. created the genetic mutants and E.R.S. performed the animal experiments. R.R.J., A.B.B., E.R.S. and J.B. analysed the data. R.R.J. and A.B.B. wrote the manuscript. All authors read and reviewed the manuscript.

