## Supplemental_FigTab_6of9 for "Prophages and satellite prophages are widespread among *Streptococcus* species and may play a role in pneumococcal pathogenesis"

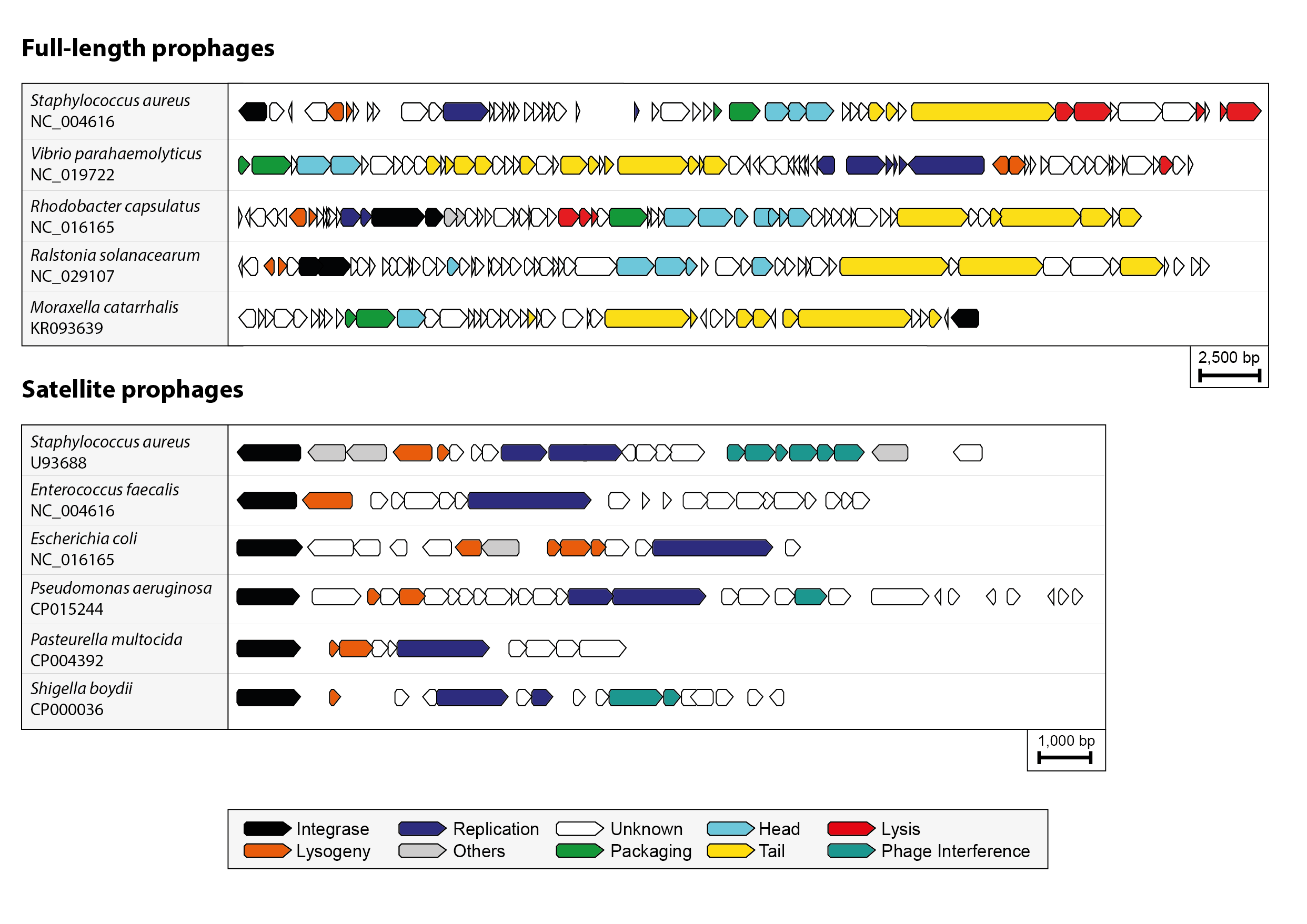

**Supplementary Figure 1.** Full-length and satellite prophages from different bacterial species demonstrate similar patterns of genome organisation and synteny.

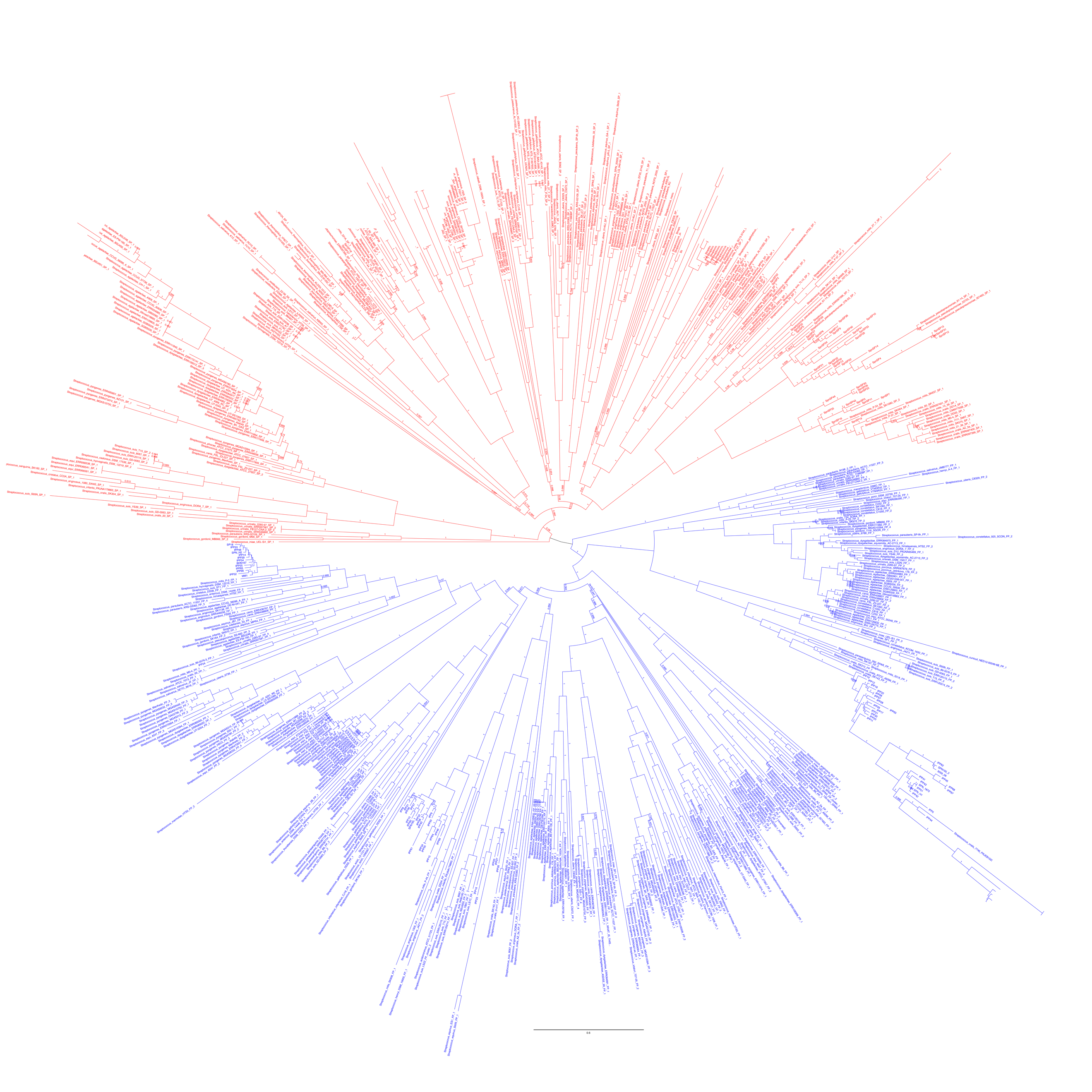

**Supplementary Figure 2.** An unrooted phylogenetic tree of all streptococcal prophage genomes identified in the dataset. Blue branches mark full-length prophages and red branches mark satellite prophages.

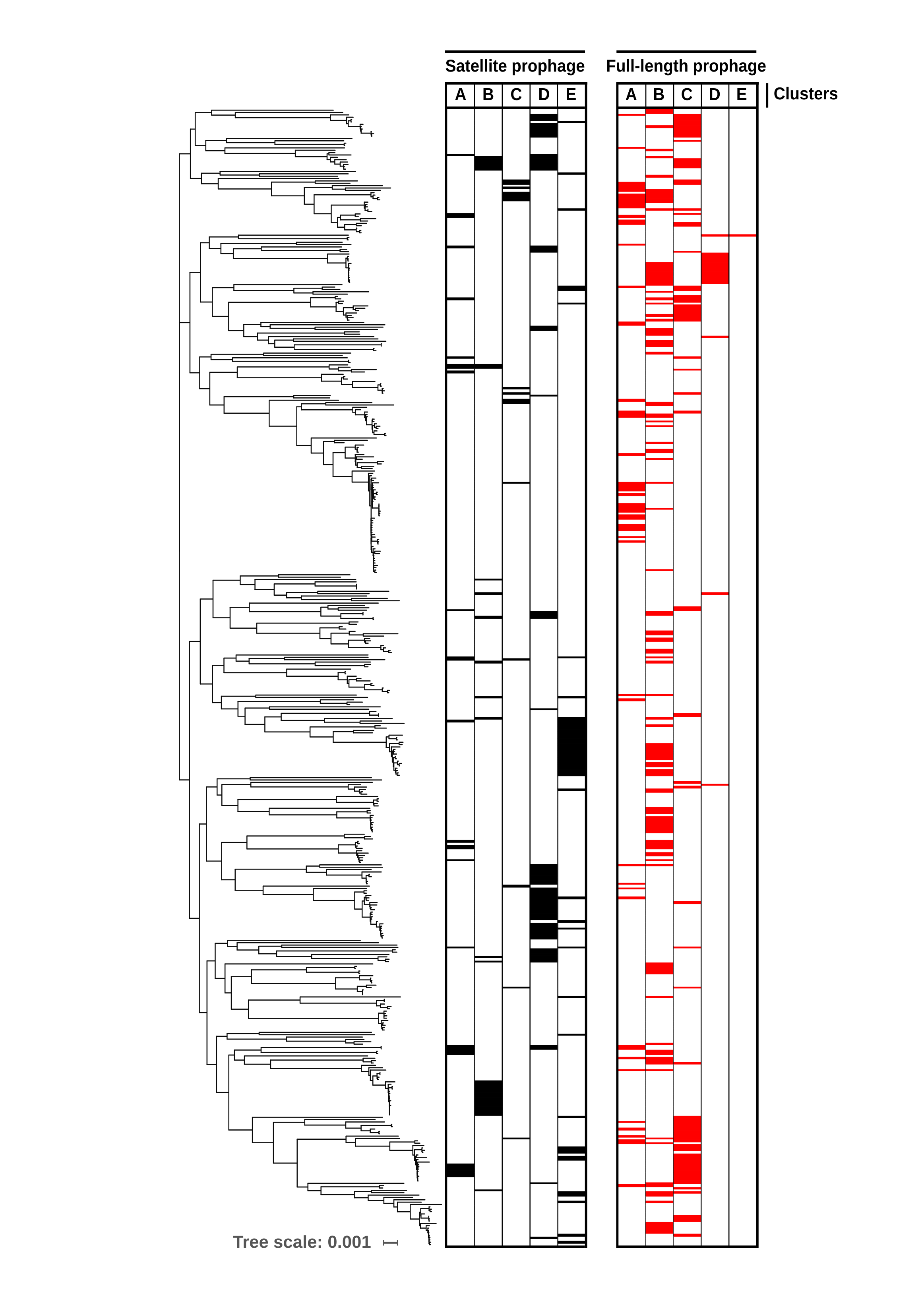

**Supplementary Figure 3.** A pneumococcal core genome phylogenetic tree annotated with corresponding satellite and full-length prophage clusters.

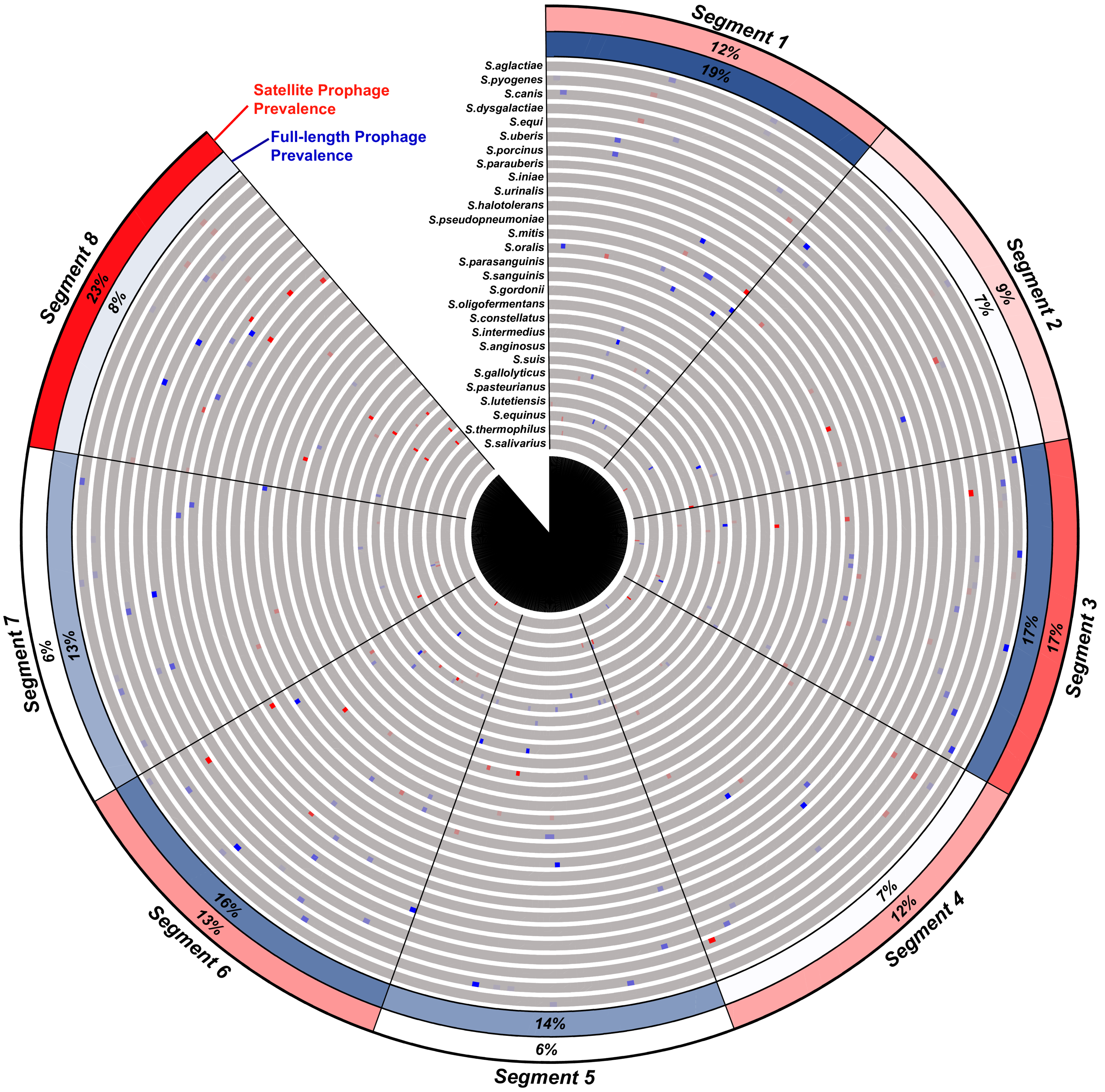

**Supplementary Figure 4.** The location of prophage insertion sites within the bacterial genomes, relative to 8 equal segments of each genome.

**Supplementary Table 1.** Descriptive statistics for full-length prophages and satellite prophages identified among 1,306 streptococcal genomes.

| **Host species** | | | **Full-length prophages** | | | | **Satellite prophages** | | | |
| --- | --- | --- | --- | --- | --- | --- | --- | --- | --- | --- |
| Name | Genomes (n) | Avg. GC content | Unique phages (n) | Avg. size (bp) | Avg. GC content | Avg. no. of genes | Unique phages (n) | Avg. size (bp) | Avg. GC content | Avg. no. of genes |
| *S. pneumoniae* | 482 | 39.6% | 66 | 37,346 | 39.6% | 56 | 44 | 12,936 | 37.2% | 21 |
| *S. pyogenes* | 50 | 38.4% | 67 | 39,113 | 38.5% | 56 | 21 | 12,825 | 36.3% | 23 |
| *S. agalactiae* | 50 | 35.3% | 38 | 37,419 | 39.3% | 51 | 19 | 14,096 | 35.7% | 23 |
| *S. dysgalactiae* | 50 | 39.3% | 35 | 39,402 | 38.6% | 57 | 24 | 13,422 | 35.6% | 24 |
| *S. suis* | 50 | 41.2% | 29 | 35,046 | 41.0% | 52 | 24 | 10,216 | 38.1% | 17 |
| *S. equi* | 50 | 41.6% | 18 | 38,096 | 39.0% | 51 | 4 | 12,813 | 36.3% | 20 |
| *S. parauberis* | 16 | 35.5% | 14 | 38,455 | 35.8% | 55 | 17 | 13,564 | 32.7% | 20 |
| *S. mitis* | 49 | 40.1% | 12 | 37,559 | 40.0% | 55 | 36 | 12,082 | 36.1% | 18 |
| *S. oralis* | 49 | 41.0% | 12 | 35,654 | 39.7% | 50 | 15 | 11,649 | 36.0% | 17 |
| *S. anginosus* | 24 | 38.6% | 10 | 36,885 | 40.1% | 49 | 17 | 11,072 | 36.7% | 17 |
| *S. equinus* | 27 | 37.3% | 10 | 40,614 | 39.1% | 56 | 13 | 10,075 | 34.2% | 15 |
| *S. constellatus* | 10 | 38.0% | 10 | 43,497 | 39.9% | 51 | 4 | 10,347 | 35.3% | 21 |
| *S. uberis* | 13 | 36.5% | 9 | 39,936 | 37.7% | 54 | 11 | 11,652 | 33.2% | 18 |
| *S. gallolyticus* | 17 | 37.5% | 8 | 37,207 | 38.1% | 52 | 9 | 9,759 | 35.1% | 14 |
| *S. canis* | 11 | 39.5% | 8 | 37,776 | 40.4% | 51 | 3 | 13,895 | 36.3% | 25 |
| *S. urinalis* | 4 | 34.1% | 7 | 38,072 | 37.2% | 53 | 8 | 9,148 | 33.8% | 14 |
| *S. parasanguinis* | 31 | 41.8% | 7 | 38,470 | 41.4% | 54 | 6 | 9,435 | 34.4% | 11 |
| *S. gordonii* | 22 | 40.4% | 7 | 35,522 | 40.1% | 47 | 3 | 10,069 | 35.4% | 15 |
| *S. pseudopneumoniae* | 16 | 39.8% | 5 | 36,007 | 39.7% | 62 | 9 | 11,870 | 38.2% | 19 |
| *S. iniae* | 8 | 36.6% | 5 | 35,285 | 36.8% | 52 | 1 | 12,203 | 30.0% | 18 |
| *S. salivarius* | 32 | 39.7% | 4 | 40,116 | 42.2% | 42 | 4 | 9,577 | 37.7% | 15 |
| *S. infantis* | 4 | 39.4% | 4 | 35,734 | 39.3% | 49 | 1 | 11,158 | 38.0% | 14 |
| *S. porcinus* | 2 | 36.7% | 4 | 37,741 | 39.6% | 51 | 0 | - | - | - |
| *S. pseudoporcinus* | 5 | 37.2% | 3 | 38,992 | 38.3% | 62 | 2 | 8,343 | 33.8% | 19 |
| *S. entericus* | 1 | 44.6% | 2 | 41,400 | 43.0% | 59 | 2 | 10,349 | 42.4% | 18 |
| *S. himalayensis* | 1 | 41.3% | 2 | 39,219 | 40.8% | 49 | 2 | 11,811 | 38.8% | 16 |
| *S. marmotae* | 1 | 40.9% | 2 | 37,995 | 43.0% | 66 | 2 | 14,088 | 40.6% | 21 |
| *S. infantarius* | 2 | 37.6% | 2 | 33,058 | 38.4% | 48 | 1 | 9,627 | 35.7% | 14 |
| *S. azizii* | 3 | 42.7% | 2 | 43,572 | 41.2% | 58 | 0 | - | - | - |
| *S. henryi* | 2 | 38.6% | 2 | 40,013 | 39.8% | 58 | 0 | - | - | - |
| *S. ictaluri* | 2 | 38.1% | 2 | 24,012 | 38.4% | 32 | 0 | - | - | - |
| *S. intermedius* | 9 | 37.6% | 1 | 33,366 | 38.0% | 49 | 6 | 13,935 | 36.4% | 23 |
| *S. pasteurianus* | 5 | 37.3% | 1 | 35,546 | 38.0% | 45 | 4 | 10,524 | 35.0% | 15 |
| *S. lutetiensis* | 2 | 37.6% | 1 | 37,997 | 39.0% | 51 | 3 | 10,881 | 35.1% | 15 |
| *S. halotolerans* | 1 | 39.2% | 1 | 41,477 | 38.1% | 52 | 2 | 13,216 | 36.9% | 20 |
| *S. hyovaginalis* | 1 | 39.9% | 1 | 39,028 | 37.6% | 60 | 2 | 12,180 | 37.0% | 22 |
| *S. acidominimus* | 1 | 42.6% | 1 | 38,191 | 40.1% | 52 | 1 | 9,826 | 39.5% | 15 |
| *S. cristatus* | 4 | 42.6% | 1 | 38,959 | 39.8% | 52 | 1 | 13,630 | 40.4% | 17 |
| *S. macedonicus* | 2 | 37.5% | 1 | 38,767 | 38.5% | 52 | 1 | 10,632 | 33.8% | 14 |
| *S. phocae* | 2 | 39.5% | 1 | 46,987 | 37.4% | 76 | 1 | 12,626 | 36.2% | 20 |
| *S. cuniculi* | 1 | 43.4% | 1 | 28,958 | 40.4% | 44 | 0 | - | - | - |
| *S. orisratti* | 1 | 38.5% | 1 | 37,447 | 41.2% | 42 | 0 | - | - | - |
| *S. porci* | 1 | 40.8% | 1 | 40,394 | 35.3% | 45 | 0 | - | - | - |
| *S. thoraltensis* | 1 | 38.4% | 1 | 40,339 | 37.5% | 57 | 0 | - | - | - |
| *S. thermophilus* | 32 | 39.0% | 0 | - | - | - | 16 | 8,429 | 37.3% | 14 |
| *S. sanguinis* | 37 | 43.0% | 0 | - | - | - | 6 | 11,336 | 38.5% | 18 |
| *S. vestibularis* | 6 | 39.5% | 0 | - | - | - | 3 | 10,768 | 36.5% | 14 |
| *S. castoreus* | 1 | 37.8% | 0 | - | - | - | 2 | 10,268 | 36.1% | 18 |
| *S. merionis* | 1 | 41.7% | 0 | - | - | - | 1 | 10,429 | 39.5% | 17 |
| *S. oligofermentans* | 1 | 42.1% | 0 | - | - | - | 1 | 10,319 | 36.2% | 16 |
| *S. caballi* | 1 | 40.4% | 0 | - | - | - | 1 | 9,970 | 38.7% | 17 |
| *S. plurextorum* | 1 | 41.1% | 0 | - | - | - | 1 | 13,633 | 35.0% | 27 |
| Other *strep.* species | 111 | - | 0 | - | - | - | 0 | - | - | - |
| **Sum** | **1,306** | **-** | **419** | **-** | **-** | **-** | **354** | **-** | **-** | **-** |
| **Average** | **-** | **39.4%** | **-** | **37,879** | **39.2%** | **53** | **-** | **11,379** | **36.2%** | **18** |

**Supplementary Table 5.** Description of the plasmid, bacterial strain and PCR primers used in the mouse experiments.

| **Name** | | **Description (source/reference/sequence)** |
| --- | --- | --- |
| **Plasmid** | pR412 | Derived from ColE1, carrying a 1145 bp minitransposon that contains Himar1 IRs flanking *addA9* gene. Spc^R^ (Martin, B., Prudhomme, M., Alloing, G., Granadel, C. y Claverys, J.P.2000) |
| **Strain** | 6B(BHN418) | *S. pneumoniae* capsular serotype 6 |
| **Primers** | SpnSP_UpF | GGTTTTCATGATGTTGTTCTGG |
|  | SpnSP_Upspec_F | GAGAGAAAAACTTGTTTCATGATCCCCCGTTTGATTTTTAAT |
|  | SpnSP_UpspecR | TTAAAAATCAAACGGGGGATCATGAAACAAGTTTTTCTCT |
|  | SpnSP_Downspec_F | ATTGGATCCATTCCGCGTCTTTAACCTTACCACGGAATTA |
|  | SpnSP_Downspec_R | TAATTCCGTGGTAAGGTTAAAGACGCGGAATGGATCCAAT |
|  | SpnSP_DownR | GATAGCTCCATGTCCGTTGATAC |
|  | SpnSP_Up confirmation | GAAAAGACCATGGTTGGGAT |
|  | SpnSP_Down confirmation | GTTCAGGTAACTCCAAAACC |
|  | VapE_UpF | GTTCCTGAAGGGGCAGATATTG |
|  | VapE_UpspecF | TAAAAAGTGAAATCTTGGAGGTAGATCCCCCGTTTGATTTTT |
|  | VapE_UpspcR | AAAAATCAAACGGGGGATCTACCTCCAAGATTTCACTTTTTA |
|  | VapE_DownspecF | TTGGATCCATTCCGCGTCGTGTGACGTTCTTTTTTT |
|  | VapE_Downspec R | CAAAAAAAGAACGTCACACGACGCGGAATGGATCCAA |
|  | VapE_DownR | CTGTATAATACCAATACGATAGCC |
|  | VapE Up confirmation | GACATCTGGAAGTTTTTGGGG |
|  | VapE Down confirmation | CCTGGTTGTTTGTGGTAGTCT |
